# Generation of multi-lineage kidney assembloids with integration between nephrons and a single exiting collecting duct

**DOI:** 10.1101/2025.02.27.640561

**Authors:** Sean B. Wilson, Inês P. Santos, Louise Wildfang, Kristiana Imsa, Melissa H. Little

**Affiliations:** Novo Nordisk Foundation Centre for Stem Cell Medicine, Faculty of Health and Medical Sciences, University of Copenhagen, DK-2200 Copenhagen, Denmark; Novo Nordisk Foundation Centre for Stem Cell Medicine, Murdoch Children’s Research Institute, Parkville, Melbourne, 3052, Australia; Department of Paediatrics, The University of Melbourne, VIC, Australia

**Keywords:** directed differentiation, kidney organoid, assembloid, pluripotent stem cell, lineage identification

## Abstract

A functional kidney requires a patent connection between all nephrons and the collecting duct network to ensure an exit path for the urinary filtrate. While it is now possible to separately direct the differentiation of pluripotent stem cells to nephron-forming or nephric duct-forming populations, the controlled integration of these two systems has not been demonstrated in human. We report the formation of an integrated kidney assembloid via the co-culture of distinct nephric duct progenitor and nephron-forming cultures in the absence of any requirement for cell enrichment. Nephrons within these kidney assembloids connect to a single exiting nephric duct with this connection forming with the early distal nephron prior to nephron segmentation. Using constitutional reporter lines, we show that the nephrons arise from the nephric progenitors and fuse with the nephric duct while the nephric duct progenitors form a single common duct without any branching. The stromal component of the nephric duct progenitor population also contributes to the medullary stroma surrounding the nephrons. Such modular assembloids address the challenge of nephron unification in an engineered kidney tissue model.

## Introduction

The human kidney is comprised of approximately 1 million segmented nephrons connected into an integrated collecting duct network through which the urinary filtrate reaches the ureter to pass to the bladder^1^. This collecting duct epithelial tree forms via repeated dichotomous branching of a side branch of the nephric duct called the ureteric bud^2^. Signals from the tip of this ureteric tree both induce nephrons to form and support the survival of nephron progenitor cells (NPCs). The distal end of each newly formed nephron invades the adjacent ureteric epithelium (UE) to make a patent connection^3,4^. In this way, all nephrons drain through the same plumbing.

It is now possible to recreate human kidney cell types via the directed differentiation of human pluripotent stem cells (hPSC)^5^ (reviewed in Little and Combes^6^). However, the generation of a functional replacement organ for the treatment of renal failure continues to face many challenges, the most significant of these being the generation of an integrated organ in which all nephrons connect to a common urinary tract. Taguchi et al.^7^ proposed that it would require the separate patterning of hPSC to anterior intermediate mesoderm (AIM) and posterior intermediate mesoderm (PIM) to provide the nephric duct (ND) and NPCs respectively. This theory derives from the knowledge that the UE arises as a side branch of the ND, itself a derivative of the AIM. Takasato et al.^5^ proposed that the anterior or posterior identity could be conferred simply by altering the initial duration of mesoderm initiation. Isolated GATA3^+^ distal nephron epithelium from kidney organoids were able to be matured to a ureteric identity^8^, illustrating the considerable plasticity of this epithelial state. This induced UE was able to connect to hPSC-derived nephrons *in vitro* after dissociation and reaggregation^8^. Uchimura et al.^9^ separately patterned to AIM and PIM and then mixed suspensions of these cultures together, reporting the presence of a UE linked to nephrons which was able to respond to aldosterone and vasopressin to express AQP2^+^, indicative of principal cell maturation. This was again an uncontrolled structure without a unified urinary exit path. It was also not possible to ascertain the origin of the cells with collecting duct identity in the final organoid.

Numerous groups have now sought to separately generate ND progenitors from which they can form a branching epithelial structure ^7,8,10–12^. Several groups have shown an ability within Matrigel to have such structures form branching epithelium expressing GATA3, SOX9 and in some instances RET when cultured in media conditions appropriate for supporting ureteric progenitors^7,8,12,13^. Taguchi et al.^7^ have demonstrated the ability to unite ureteric, nephron and stromal progenitor populations to form impressive integrated nephrons and UE referred to as higher order kidneys^7,13^. However, this was only possible using murine and not human PSC. Shi et al.^14^ recently reported the combination of ND progenitors and dissociated NPCs showing a capacity for the formation of contiguous nephrons and UE in an organoid setting. However, this did not provide a controlled integration of nephrons with a single ureteric exit path which is the essential architecture of the organ itself.

In this study, we describe an approach to combining hPSC-derived NPCs with ND progenitors that generate an assembloid comprising a cohesive network of nephrons with a single urinary exit. Via aggregation of a single ND progenitor aggregate with NPCs in a micropatterned well format, a unified organoid in which patterned nephrons connect to this single nephric duct was generated, resulting in unified plumbing of the tissue. Distal nephron EPCAM^+^CDH1^+^ epithelium is contiguous with an EPCAM^+^ CDH1^+^ epithelium arising from the ND progenitors. Using constitutional and promoter-specific fluorescent reporter hPSC lines we show the origin of this contiguous epithelium demonstrating fusion of nephrons to the ND without ND branching. This outcome has implications for the engineering of replacement renal tissue.

## Results

### Upscaling nephric duct culture for use in assembloid cultures

Using previously established protocols for the generation of both the ND^12^ andNPC^5^ derived lineages of the kidney from hPSC, we sought to combine these two progenitor populations together in an assembloid manner to better facilitate connection to a single exit path for urinary filtrate. To enable scaling of this process, the ND protocol was adapted by seeding between 3×10^4^ and 1.2×10^5^ cells at day 3 stage into single 24-well AggreWell400 plates, forming ∼1200 spheroids per well with an average of 250, 500, 750 or 1000 cells each. Using a GATA3^mCherry^ iPSC reporter line^15^, all spheroids were confirmed to be GATA3^+^ at day 7 with a small population of GATA3^-^ cells, as originally reported^12^ (Figure 1A). These structures generate CDH1^+^ epithelial structures with RET^+^ tip regions when embedded in Matrigel, as anticipated^12^ (Figure 1B). Larger spheroid wells became acidic quite quickly due to the higher metabolic output from increased overall cell numbers. Hence, seeding densities of 3-6×10^5^ cells were used in assembloid cultures.

**Figure 1:**
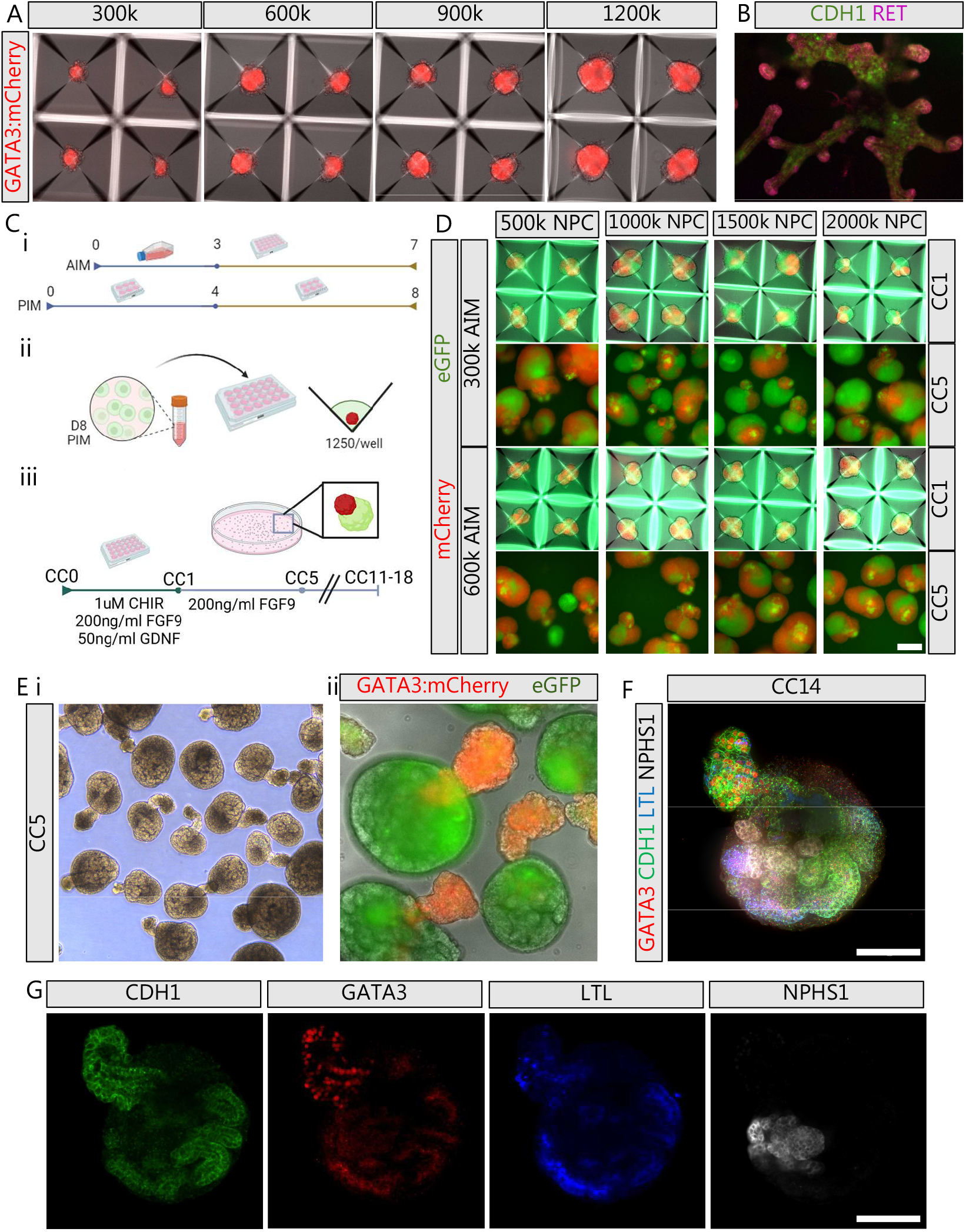
Generation of multi-lineage kidney assembloids. A) Spheroids generated by adding 3, 6, 9 or 12 x 10^5^ cells at day 3 of ND culture, the majority of cells showing positive GATA3:mCherry expression. B) Spheroids embedded in Matrigel and matured using ureteric epithelial supporting growth factors, showing branching into CDH1^+^ epithelial structures tipped with RET^+^ at day 14 of culture. C) Graphical representation of the protocol. i) The two lineages are generated in parallel following the 7 day ND protocol^12^ and the 8 day NPC protocol^5,22^, reaching days 7 and 8 respectively on the same day. ii) At this stage, NPC cells are dissociated and added to the ND spheroids in AggreWell400 plates. iii) After 1 day of culture, the assembloids are transferred to suspension culture until an end-point is reached. D) Assembloids after 5 days of assembly culture generated from either 3 or 6 x 10^5^ day 3 ND spheroids (red) grown to day 7 and 5, 10, 15 or 20 x 10^5^ day 8 NPC cells (green). E) Assembloid morphology i) showing the epithelial extension by 5 days of co-culture ii) day 14 assembloids showing dysmorphic morphology in the ureteric epithelium present in many late-stage assembloids. F) and G) Segmentation analysis of day 14 assembloids shows the extension is CDH1^+^/GATA3^+^, connected to the nephron network with segmentation from the CDH1^+^ distal tubule, LTL^+^ proximal tubule and NPHS1^+^ podocytes in the glomerulus. Scale bar 100µm; k, thousand; ND, nephric duct; NPC, nephron progenitor cell; CC, co-culture

To generate an assembloid, constitutionally GFP-labelled cells from a nephron progenitor differentiation^5^ were dissociated and added to day 7 nephric duct progenitor spheroids generated using the GATA3^mCherry^ line (Figure 1C). This enabled an investigation of the resulting morphology and proportion of each line in the co-culture. 5, 10, 15 or 20 x 10^5^ dissociated NPCs were plated per single well of a 24-well AggreWell400, with each well containing single spheroids in microwells (Figure 1Cii). After 24 hours, the aggregated assembloids were transferred to suspension culture for an additional 4 days (Figure 1Ciii). The proportion of each cell source within assembloids was evaluated at this point using the reporter expression (Figure 1D). Assembloids comprised of 3×10^5^ nephric duct progenitor spheroid and 10 or 15×10^5^ NPC showed the most optimal cell type ratio and morphology, comprising a GFP-positive main body with a distinct RFP-positive region.

### Assembloid co-culture generates kidney organoids with connected ureteric and nephron epithelia

Following 5 days of co-culture (CC5), a single noticeable cellular extension perpendicular to the main assembloid body was present (Figure 1Ei). This extension was epithelial and contained only mCherry+ ND-lineage cells (Figure 1Eii). After 14 days of co-culture (CC14), assembloids displayed a network of epithelium comprised of segmented nephrons (Figure 1F), with evidence of CDH1^+^/GATA3^+^ distal tubule, LTL^+^ proximal tubules and NPHS1^+^ podocytes (Figure 1G). The extension was CDH1^+^/GATA3^+^. This epithelial structure was directly connected to the forming and segmenting nephrons within the main body of the assembloid (Figure 1F, 1G). As assembloids develop beyond 11 days in suspension culture the ureteric structure became dysmorphic (Figure 1Eii) with some detaching from the main body of the assembloid. This indicates a sub-optimal culture environment for this structure.

This protocol leads to the development of assembloids containing a single ureteric structure, a feature lacking from existing human kidney organoid protocols. Establishing this culture in AggreWell400 24-well plates allowed for the generation of individual assembloids within each of the 1200 microwells. A single confluent T25 flask differentiation of each lineage is projected to generate ∼9000 independent assembloids at a time.

### Constitutional reporter lines confirm epithelial connection between nephrons and a single ureteric structure

Constitutive fluorescent human iPSC lines were used to generate the ND (EEF2^mCherry^, unpublished) and NPC (GAPDH^eGFP^) ^16^ differentiations. This enabled tracing of the lineage origin of all cells in the final assembloid, visualization of the epithelial networks formed in the assembloid and investigation of fusion between lineages. After 24 hours of co-culture in an AggreWell 400 plate, the resulting assembloids contained a single mCherry^+^ epithelial structure (Figure 2Ai). These were transferred to suspension for ongoing culture following our previous protocol^17^ (Figure 2Aii). Following a further 4 days of culture, the cellular extension adjacent to the main body of each assembloid was mCherry^+^ and contained no eGFP^+^ cells, indicating an ND lineage origin (Figure 2Aiii). By day 5, the NPC population generated an EPCAM^+^ structure adjacent to the ND-derived epithelium connecting to the ND-derived epithelium (Figure 2B). At high resolution, the mCherry^+^ is seen to be contiguous with the mCherry^-^ epithelium (Figure 2C). Epithelial structures comprised of both component populations can be identified at this time point.

**Figure 2:**
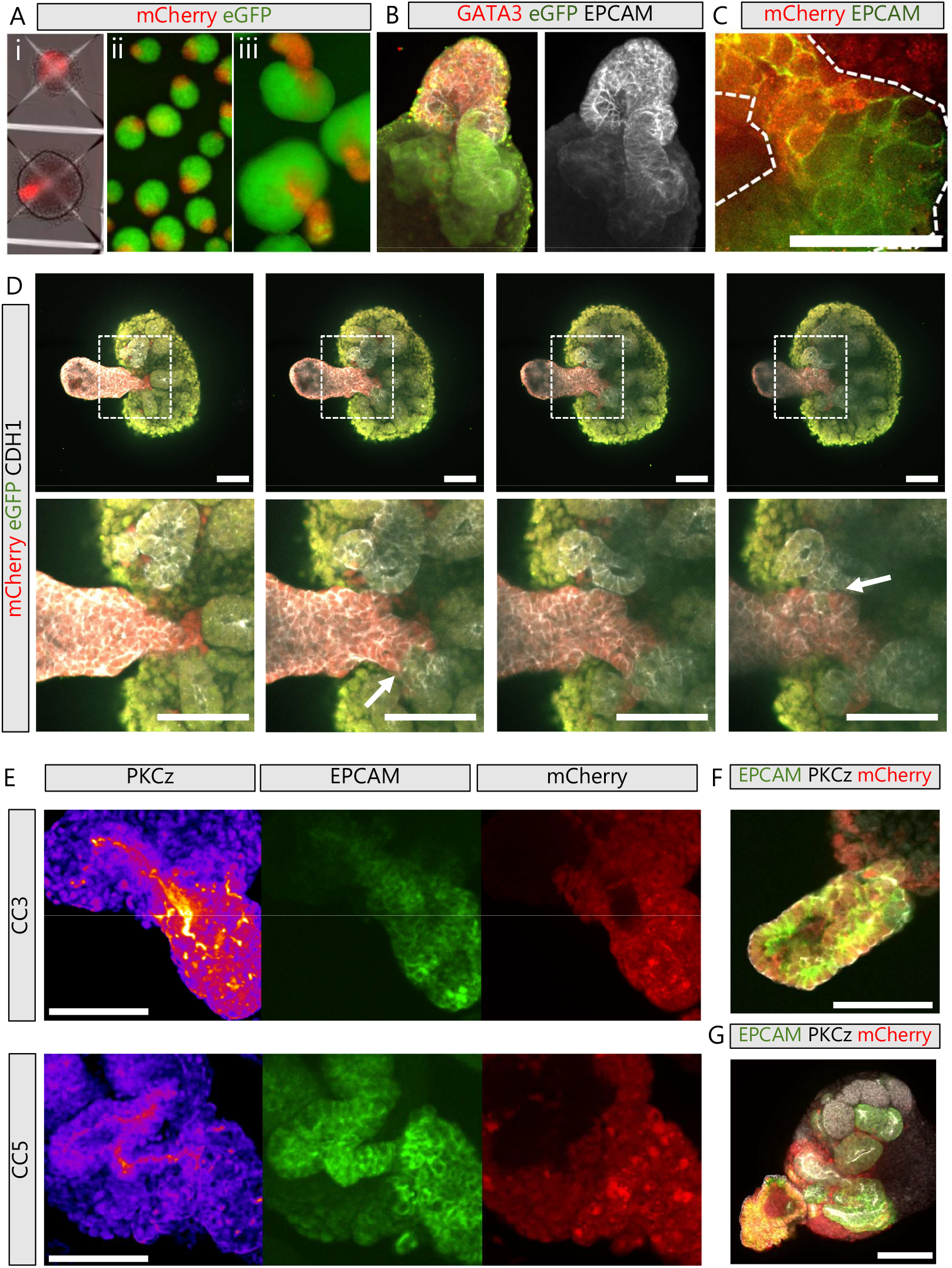
Assembloid culture leads to the generation of a connected epithelial network. A) Assembloids after 1 day (i) in the AggreWell400 plate or (ii) after transfer to suspension culture shows the ND-derived (mCherry) spheroid surrounded by NPC-derived mesenchyme, which by day 5 (iii) have formed a structure extending away from the main body of the assembloid. B) Day 5 assembloids contain an NPC-derived (GFP) EPCAM^+^ epithelium adjacent to an ND-derived (GFP^-^) EPCAM^+^/GATA3^+^ epithelium. C) An epithelial structure (white dotted line) in a day 5 assembloid showing the fusion between ND-derived (mCherry^+^) and NPC-derived (mCherry^-^) epithelium. Scale bar is 50µm D) Serial images through an 8 day post co-culture (CC8) assembloid. Dotted box expanded in the second row is the connection points between the NPC-derived nephrons (CDH1^+^/GFP^+^) to the ND-derived ureteric epithelium (CDH1^+^/mCherry^+^). The white arrows show the two nephron-ureteric connection points. Scale bar is 100µm. E) Maximum intensity projection of a 3 day (CC3, top) and 5 day (CC5, bottom) assembloid showing the apical membrane (PKCz) of a connected EPCAM+ epithelium fused between the mCherry^+^ and mCherry^-^ lineages. Scale bar is 100µm. F) The inverted polarization of a 3 day assembloid ND-derived epithelial extension, with apical marker PKCz on the external surface of the epithelium. Scale bar is 100µm. G) The 14 day assembloid shows a dysmorphic epithelial extension yet appropriately polarized nephrons within the assembloid main body. Scale bar is 100µm. NPC, nephron progenitor cell; ND, nephric duct; CC, coculture day.

At 8 days of co-culture (CC8) the epithelial network has advanced and become more complex. The connection between the CDH1^+^/RFP^+^ and CDH1^+^/GFP^+^ epithelium is clearly identifiable, further demonstrating a connection between epithelium from both lineages (Figure 2D). Interestingly, it appeared that multiple nephrons can form connection points to the UE (Figure 2D, arrows). Investigation of luminal connection was performed using atypical protein kinase C zeta (PKCz) to mark the apical lumen at three and five days of assembloid co-culture (CC3/CC5). At CC3 a single continuous epithelium with PKCz was identified across the fused epithelium (Figure 2E). At CC5, the epithelium had become more extensive with an expanded NPC-derived epithelium present. This epithelium exhibited a single PKCz lumen that remained fused between the two lineages (Figure 2E). However, the majority of ND-derived epithelium forming the extension away from the main assembloid body did not contain an organized PKCz lumen structure, despite being EPCAM expressing. Instead, the extension becomes inverted, showing PKCz expression on the external, media facing surface (Figure 2F). This leads to the dysmorphic structure present in mature assembloids (Figure 2F).

This assembly method for co-culture of the ND and NPC populations leads to a robust generation of assembloids containing a single ND-derived ureteric epithelium connected to NPC-derived nephrons, thereby forming a single path of exit for urinary filtrate (Figure 2E). We confirmed fusion in 85.7% of cases (7 biological replicates), with these connections occurring via fusion or integration of the NPC-derived epithelium into the single ND-derived epithelium.

### Interaction between the two lineages creates a nephrogenic niche environment and connected epithelium but no ureteric branching

*In vivo,* the reciprocal interaction between the ureteric tip and surrounding nephron progenitors drives branching morphogenesis and ongoing nephron commitment^2^. To answer whether anticipated signalling between these two populations was evident, immunofluorescent staining of RET and SIX2, markers of the UE tip and NPC respectively, were investigated.

RET is expressed in the ND-lineage after 1 day of co-culture (CC1), localized to the border adjacent to NPC-lineage cells (Figure 3A). By 3 days of co-culture (CC3), the epithelial fusion between lineages was evident, with RET expression present on the ND-derived cells at that border (Figure 3B). Concurrently, SIX2^+^ cells derived from the NPC-lineage (mCherry^-^) are present (Figure 3C). The adjacency of both ureteric tip and nephrogenic mesenchyme is reminiscent of the *in vivo* nephrogenic niche; however, the characteristic budding of the tip was not seen. *In vivo,* the ND extends posteriorly with mesenchymal cell protrusions into adjacent space^18^. This same behaviour is seen in the assembloids (Figure 3D). By 5 days of co-culture (CC5) SIX2^+^ cells surrounded the growing epithelial structures. At this time point it was much rarer to see RET^+^ ureteric tips with no evidence of branching within the ND lineage (Figure 3E). The lack of bifurcating ureteric tips and capacity for ND-like migrational growth indicates the appropriate reciprocal induction was not initiated in the *in vitro* niche to fully commit the RET^+^ cells to an *in vivo-*like tip domain.

**Figure 3:**
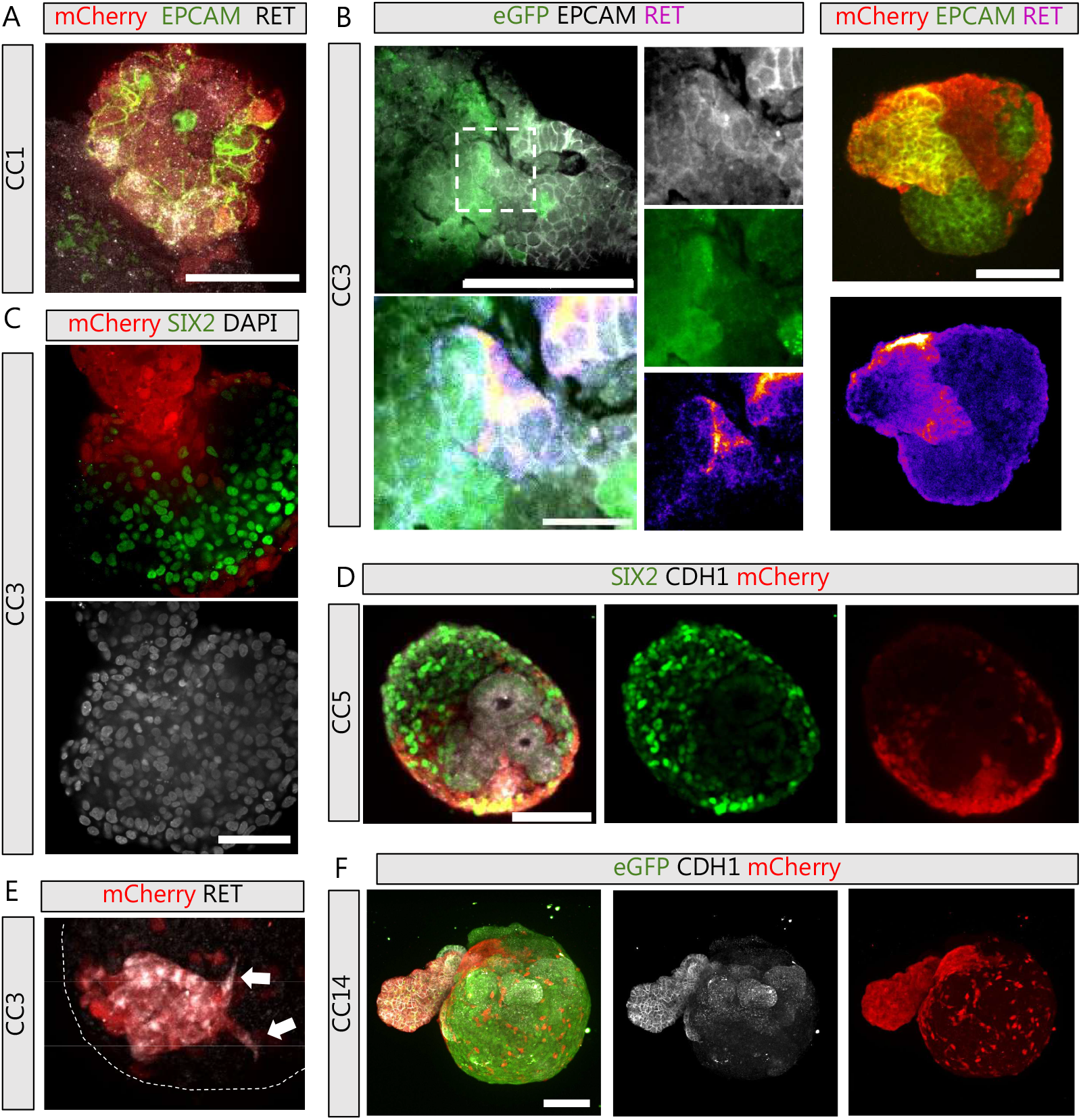
Evidence of nephrogenic niche generation and medullary mesenchymal migration. A) Expression of RET is identified by 1 day assembloid (CC1) in the ND-derived cells (mCherry^+^). Scale bar 50µm. B) Expression of RET localized to the ND-derived (GFP^-^ or mCherry^+^) epithelium after 3 days of culture (CC3), bordering the NPC-derived (GFP^+^ or mCherry^-^) mesenchyme. Scale bar 100µm (top left), 20µm (bottom left), 50µm (top right). C) Expression of SIX2 can be seen within the main body of day 3 assembloids (CC3) adjacent to the ND-derived population. Scale bar 50µm. D) Assembloids at day 5 (CC5) contain fused multi-lineage epithelium surrounded by SIX2^+^ cells. Scale bar 100µm E) Evidence of ND-derived RET^+^ cells with a extension migration morphology. F) Maximum intensity projection of a single assembloid showing the extent of ND-derived (mCherry^+^) mesenchymal cell migration thoughout the main assembloid body surrounding the developed nephrons. A concentration of these cells also remains in the medullary region nearest the ND-derived epithelium. Scale bar 100µm. CC, coculture day; ND, nephric duct;

The ND-derived component contains both a GATA3^+^ and GATA3^-^ population, with the GATA3^-^ population being a stromal subset that does not epithelialise^12^. These cells migrate throughout the nephron-containing region of the assembloid, while a population of these cells remain near the ND-derived structure (Figure 3F). As in existing organoid cultures, the ratio of mesenchymal cell to epithelial tubules between batches shows considerable technical variability, but always results in a significant proportion of mesenchyme within the final organoid (Figure 3F). The exact identity of these cells beyond mesenchymal was not further investigated.

## Conclusions

The generation of assembloid kidney organoids addresses key challenges in the field. Separately generating each lineage and adding the nephron generating cells onto a single ureteric precursor spheroid generated a contiguous tubular network with a single unified exit path. However, the formation of a connection between nephric duct and nephrons did not resemble the process observed in metanephric development *in vivo.* The interaction between these two populations did not lead to ureteric branching, rather the nephrons induced to form immediately connected with the adjacent UE. This may be more reminiscent of mesonephric tubule fusion to the nephric duct^19^. This suggests the lack of appropriate niche components able to support epithelial branching. Conversely, the induction of nephrons prior to the formation of a ureteric tip may also drive fusion without branching.

As assembloid culture continued, the ureteric extension that directly interfaced with the media in suspension displayed a disorganized epithelial identity, showing an inverted polarity lacking a single clear lumen rather than forming a tight extending ureter. This aberrant polarization and abnormal morphology only occurred on the media-facing section of the ND-derived epithelium and may have been exacerbated by sheer stress in the culture. Currently, the only established method of differentiation and maturation of the collecting duct *in vitro* requires Matrigel embedding. This suggests the need for an extracellular matrix or similar support.

In comparison to all previous reports in which there has been observed connections with nephrons formed between human pluripotent stem cell-derived nephric duct, ureteric epithelium or anterior intermediate mesoderm, this is the first example of an approach in which nephrons connect with a single exit path. Shi et al (2024) have also shown an ability for this nephric duct protocol to form connections, but this was not in an organized fashion with a single exiting epithelial duct. It was also not amenable to scale out as has been demonstrated here. Using this approach provides a capacity to generate comprehensive modular nephron-ureteric units at scale which may provide a considerable advantage in tissue engineering.

This current assembloid method does not allow for ongoing nephron induction due to a lack of ureteric branching and hence no ongoing support for nephron progenitor self-renewal. A single ureter with so few connected nephrons will not provide sufficient renal function. However, the kidney is a modular organ and this organ in larger mammals contains repeating ‘lobes’ of smaller connected regions of nephrons. The human kidney has 7-18 lobes while whale kidney contains over 100 lobes called ‘reniculi’ organized in a similar manner^20^. It may ultimately be possible to engineer a larger modular tissue from multiple assembloids.

In summary, we have developed a robust protocol for the assembloid co-culture of ureteric and nephron lineages to form an organoid with a connected and segmented collecting duct-nephron epithelial network. We demonstrate the capacity for the scale-out of this method to easily generate 1000s of small organoids in each culture. This protocol leverages existing single-lineage differentiation protocols and is amenable to high-throughput screening. As such, this assembloid approach is another step to advance kidney organoid technologies towards a cellular therapy.

## Acknowledgements

We acknowledge the MCRI gene editing facility for the provision of all reporter stem cell lines and support from the microscopy facility within reNEW Copenhagen. MHL and SBW are funded by the Novo Nordisk Foundation Center for Stem Cell Medicine (NNF21CC0073729). The composition of matter described in this manuscript has been patented by the University of Copenhagen.

## Author contributions

SBW and MHL conceived the study. SBW designed experimental procedures. SBW, IPS, LW and KI performed kidney organoid differentiations and analysis. SBW and MHL wrote the manuscript while all authors assisted in manuscript preparation.

## Declaration of interests

MHL and SBW are inventors on additional patents relating to the generation of kidney organoids.

## Methods

### Cell lines used

Two iPSC lines were utlised in this study; GAPDH:eGFP^16^ and EEF2:mCherry (unpublished). All iPSC lines were maintained and expanded at 37°C, 5% CO2, and 5% O2 in Essential 8 medium (Thermo Fisher Scientific) on Matrigel□coated plates with daily medium changes and passaged every 3–4 days with EDTA in 1X PBS as previously described^21^.

### Generation of assembloid cultures

Dissociate 3-6×10^5^ cells of anterior intermediate mesoderm cells (Shi protocol, day 3) in 2mL of Essential-6 (E6) with 50ng/mL FGF9 and 0.1uM all-trans retinoic acid. Transfer cell suspension into a 24-well AggreWell 400. After 2 days of culture, remove 1ml of media and replace with 1ml of E6 with 0.1uM all-trans retinoic acid and 50ng/ml GDNF. At day 7, dissociate 1.5-3 x 10^6^ posterior intermediate mesoderm cells (Takasato/Howden protocol, day 7-9) and resuspend in 1ml of E6 with 2uM CHIR99102, 400ng/ml FGF9, 2ug/ml Heparin and 100ng/ml GDNF. Perform a wash of the day 7 nephric duct spheroids in the AggreWell400 by removing 1ml of media and replacing with 1ml of E6 media twice, leaving 1ml of media only in the well after the second wash. Transfer the 1ml cell suspension into the well and allow to settle overnight. The next day, transfer the assembloids to a 15ml tube and remove the media. Resuspend in 5ml of E6 with 200ng/ml FGF9, 1ug/ml Heparin and 0.5x Anti-Anti, then transfer assembloid suspension to a low attachment 6cm^2^ culture dish. Place culture dish onto an orbital shaking platform at 60rpm. Replace with the same media after 2 days. After a further 2 days, replace with collecting duct maturation media of 5ml E6 with 10nM aldosterone, 10nM vasopressin and 0.5x Anti-Anti.

### Immunofluorescence assay of kidney assembloids

Kidney assembloids were fixed using 4% PFA in PBS for 20 minutes, before washing 3x in PBS and storing at 4 degrees Celsius. For immunostaining, organoids were first blocked using 10% donkey serum in PBS + 0.5% Triton-X for 2 hours at room temperature. Organoids were then washed 3x in PBS then incubated in primary or secondary antibody solution overnight at 4 degrees Celsius or 2 hours at room temperature. Following each incubation, organoids were washed 3x in PBS for 30 minutes at room temperature. Immunofluorescence staining was performed using the following antibodies: mouse anti-EPCAM-biotinylated (Thermo Fischer #13-9326-82), goat anti-RET (R&D Systems #AF1485), rabbit anti-SIX2 (Proteintech #11562-1-AP), chicken anti-GFP (Sapphire Bioscience #ab13970), rabbit anti-RFP (MBL, #PM005), goat anti-RFP (Novus Biological #NBP3-05558), mouse anti-MEIS1/2/3 (Activ Motif, #ATM39795), goat anti-GATA3 (R&D Systems, $AF2605), mouse anti-CDH1 (BD Biosciences, #610182). Organoids were imaged using a Nikon spinning disk confocal microscope.

## References

1. Bertram, J. F., Douglas-Denton, R. N., Diouf, B., Hughson, M. D. & Hoy, W. E. Human nephron number: implications for health and disease. Pediatr. Nephrol. 26, 1529 (2011).

2. Costantini, F. & Kopan, R. Patterning a Complex Organ: Branching Morphogenesis and Nephron Segmentation in Kidney Development. Dev. Cell 18, 698–712 (2010).

3. Georgas, K. et al. Analysis of early nephron patterning reveals a role for distal RV proliferation in fusion to the ureteric tip via a cap mesenchyme-derived connecting segment. 332, 273–286 (2009).

4. Kao, R. M., Vasilyev, A., Miyawaki, A., Drummond, I. A. & McMahon, A. P. Invasion of distal nephron precursors associates with tubular interconnection during nephrogenesis. J. Am. Soc. Nephrol. JASN 23, 1682–1690 (2012).

5. Takasato, M. et al. Kidney organoids from human iPS cells contain multiple lineages and model human nephrogenesis. Nature 526, 564–568 (2015).

6. Little, M. H. & Combes, A. N. Kidney organoids: accurate models or fortunate accidents. Genes Dev. 33, 1319–1345 (2019).

7. Taguchi, A. & Nishinakamura, R. Higher-Order Kidney Organogenesis from Pluripotent Stem Cells. Cell Stem Cell 21, 730-746.e6 (2017).

8. Howden, S. E. et al. Plasticity of distal nephron epithelia from human kidney organoids enables the induction of ureteric tip and stalk. Cell Stem Cell (2020) doi:10.1016/j.stem.2020.12.001.

9. Uchimura, K., Wu, H., Yoshimura, Y. & Humphreys, B. D. Human Pluripotent Stem Cell-Derived Kidney Organoids with Improved Collecting Duct Maturation and Injury Modeling. Cell Rep. 33, 108514 (2020).

10. Tanigawa, S. et al. Activin Is Superior to BMP7 for Efficient Maintenance of Human iPSC-Derived Nephron Progenitors. Stem Cell Rep. (2019) doi:10.1016/j.stemcr.2019.07.003.

11. Mae, S.-I. et al. Expansion of Human iPSC-Derived Ureteric Bud Organoids with Repeated Branching Potential. Cell Rep. 32, 107963 (2020).

12. Shi, M. et al. Human ureteric bud organoids recapitulate branching morphogenesis and differentiate into functional collecting duct cell types. Nat. Biotechnol. 41, 252–261 (2023).

13. Tanigawa, S. et al. Generation of the organotypic kidney structure by integrating pluripotent stem cell-derived renal stroma. Nat. Commun. 2022 131 13, 1–15 (2022).

14. Shi, M. et al. Integrating collecting systems in kidney organoids through fusion of distal nephron to ureteric bud. 2024.09.19.613645 Preprint at 10.1101/2024.09.19.613645 (2024).

15. Vanslambrouck, J. M. et al. A Toolbox to Characterize Human Induced Pluripotent Stem Cell-Derived Kidney Cell Types and Organoids. J. Am. Soc. Nephrol. JASN 30, 1811–1823 (2019).

16. Howden, S. E., Vanslambrouck, J. M., Wilson, S. B., Tan, K. S. & Little, M. H. Reporter-based fate mapping in human kidney organoids confirms nephron lineage relationships and reveals synchronous nephron formation. EMBO Rep. 0, e47483 (2019).

17. Kumar, S. V. et al. Kidney micro-organoids in suspension culture as a scalable source of human pluripotent stem cell-derived kidney cells. Development 146, dev172361 (2019).

18. Sanchez-Ferras, O. et al. A coordinated progression of progenitor cell states initiates urinary tract development. Nat. Commun. 12, 2627 (2021).

19. Sainio, K., Hellstedt, P., Kreidberg, J. A., Saxén, L. & Sariola, H. Differential regulation of two sets of mesonephric tubules by WT-1. Development 124, 1293–1299 (1997).

20. Ortiz, R. M. Osmoregulation in Marine Mammals. J. Exp. Biol. 204, 1831–1844 (2001).

21. Chen, G. et al. Chemically defined conditions for human iPSC derivation and culture. Nat. Methods 8, 424–429 (2011).

22. Howden, S. E. & Little, M. H. Generating Kidney Organoids from Human Pluripotent Stem Cells Using Defined Conditions. In Stem Cells and Tissue Repair □: Methods and Protocols (ed. Kioussi, C.) 183–192 (Springer US, New York, NY, 2020). doi:10.1007/978-1-0716-0655-1_15.

